# REMoDNaV: Robust Eye-Movement Classification for Dynamic Stimulation

**DOI:** 10.1101/619254

**Authors:** Asim H. Dar, Adina S. Wagner, Michael Hanke

## Abstract

Tracking of eye movements is an established measurement for many types of experimental paradigms. More complex and more prolonged visual stimuli have made algorithmic approaches to eye movement event classification the most pragmatic option. A recent analysis revealed that many current algorithms are lackluster when it comes to data from viewing dynamic stimuli such as video sequences. Here we present an event classification algorithm—built on an existing velocity-based approach—that is suitable for both static and dynamic stimulation, and is capable of classifying saccades, post-saccadic oscillations, fixations, and smooth pursuit events. We validated classification performance and robustness on three public datasets: 1) manually annotated, trial-based gaze trajectories for viewing static images, moving dots, and short video sequences, 2) lab-quality gaze recordings for a feature length movie, and 3) gaze recordings acquired under suboptimal lighting conditions inside the bore of a magnetic resonance imaging (MRI) scanner for the same full-length movie. We found that the proposed algorithm performs on par or better compared to state-of-the-art alternatives for static stimulation. Moreover, it yields eye movement events with biologically plausible characteristics on prolonged dynamic recordings. Lastly, algorithm performance is robust on data acquired under suboptimal conditions that exhibit a temporally varying noise level. These results indicate that the proposed algorithm is a robust tool with improved classification accuracy across a range of use cases. The algorithm is cross-platform compatible, implemented using the Python programming language, and readily available as free and open source software from public sources.

## Introduction

A spreading theme in cognitive neuroscience is to use dynamic and naturalistic stimuli such as video clips or movies as opposed to static and isolated stimuli (Matusz et al., 2019). Using dynamic stimuli promises to observe the nuances of cognition in a more life-like environment (Maguire, 2012). Some interesting applications include the determination of neural response to changes in facial expression (Harris et al., 2014), understanding complex social interactions by using videos (Tikka et al., 2012), and more untouched themes such as the underlying processing of music (Toiviainen et al., 2014). In such studies, an unobtrusive behavioral measurement is required to quantify the relationship between stimulus and response. Tracking the focus of participants’ gaze is a suitable, well established method that has been successfully employed in a variety of studies ranging from the understanding of visual attention (Liu and Heynderickx, 2011), memory (Hannula et al., 2010), and language comprehension (Gordon et al., 2006). Regardless of use case, the raw eye tracking data (gaze position coordinates) provided by eye tracking devices are rarely used “as is”. Instead, in order to disentangle different cognitive, oculomotor, or perceptive states associated with different types of eye movements, most research relies on the classification of eye gaze data into distinct eye movement event categories (Schutz et al., 2011). The most feasible approach for doing this lies in the application of appropriate event classification algorithms.

However, a recent comparison of algorithms found that while many readily available algorithms for eye movement classification performed well on data from static stimulation or short trial-based acquisitions with simplified moving stimuli, none worked particularly well on data from complex dynamic stimulation, such as video clips, when compared to human coders (Andersson et al., 2017). And indeed, when we evaluated an algorithm by Nyström and Holmqvist (2010), one of the winners in the aforementioned comparison, on data from prolonged stimulation (≈15 min) with a feature film, we found the average and median durations of labeled fixations to exceed literature reports (e.g., Holmqvist et al., 2011; Dorr et al., 2010) by up to a factor of two. Additionally, and in particular for increasing levels of noise in the data, the algorithm classified too few fixations, as also noted by Friedman et al. (2018), because it discarded potential fixation events that contained data artifacts such as signal-loss and distortion associated with blinks.

Therefore our objective was to improve upon the available eye movement classification algorithms, and develop a tool that performs robustly on data from dynamic, feature-rich stimulation, without sacrificing classification accuracy for static and simplified stimulation. Importantly, we aimed for applicability to prolonged recordings that potentially exhibit periods of signal-loss and non-stationary noise levels. Finally, one of our main objectives was to keep the algorithm as accessible and easily available as possible in order to ease the difficulties associated with closed-source software or non-publicly available source code of published algorithms.

Following the best practices proposed by Hessels et al. (2018), we define the different eye-movements that are supported by our algorithm on a functional and oculomotor dimension as follows: A *fixation* is a period of time during which a part of the visual stimulus is looked at and thereby projected to a relatively constant location on the retina. This type of eye movement is necessary for visual intake, and characterized by a relatively still gaze position with respect to the world (e.g., a computer screen used for stimulus presentation) in the eye-tracker signal. A fixation event therefore excludes periods of *smooth pursuit*. These events are eye movements during which a part of the visual stimulus that moves with respect to the world is looked at for visual intake (e.g., a moving dot on a computer screen). Like fixations, the stimulus is projected to a relatively constant location on the retina (Carl and Gellman, 1987), however, the event is characterized by steadily changing gaze position in the eye-tracker signal. If this type of eye movement is not properly classified, erroneous fixation and saccade events (which smooth pursuits would be classified into instead) are introduced (Andersson et al., 2017). Contemporary algorithms rarely provide this functionality (but see e.g., Larsson et al., 2015; Komogortsev and Karpov, 2013, for existing algorithms with smooth pursuit classification). *Saccades* on the other hand are also characterized by changing gaze positions, but their velocity trace is usually higher than that of pursuit movements. They serve to shift the position of the eye to a target region, and, unlike during pursuit or fixation events, visual intake is suppressed (Schutz et al., 2011). Lastly, *post-saccadic oscillations* are periods of ocular instability after a saccade (Nyström and Holmqvist, 2010).

Here we introduce REMoDNaV (robust eye movement classification for dynamic stimulation), a new tool that aims to meet our objectives and classifies the eye movement events defined above. It is built on the aforementioned algorithm by Nyström and Holmqvist (2010) (subsequently labeled NH) that employs an adaptive approach to velocity based eye movement event classification. REMoDNaV enhances NH with the use of robust statistics, and a compartmentalization of prolonged time series into short, more homogeneous segments with more uniform noise levels. Furthermore, it adds support for pursuit event classification. Just as the original algorithm, its frame of reference is world centered, i.e. the gaze coordinates have a reference to a stimulation set-up with a fixed position in the world such as x and y coordinates in pixel of a computer screen, and it is meant to be used with eye tracking data from participants viewing static (e.g. images) or dynamic (e.g. videos) stimuli, recorded with remote or tower-mounted eye trackers. Importantly, it is built and distributed as free, open source software, and can be easily obtained and executed with free tools. We evaluated REMoDNaV on three different datasets from conventional paradigms, and dynamic, feature-rich stimulation (high and lower quality), and relate its performance to the algorithm comparison by Andersson et al. (2017).

## Methods

Event classification algorithms can be broadly grouped into *velocity-* and *dispersion*-based algorithms. The former rely on velocity thresholds to differentiate between different eye movement events, while the latter classify eye movements based on the size of the region the recorded data falls into for a given amount of time (Holmqvist et al., 2011). Both types of algorithms are common (see e.g., Hessels et al. (2017) for a recent dispersion-based, and e.g., van Renswoude et al. (2018) for a recent velocity-based solution, and see Dalveren and Cagiltay (2019) for an evaluation of common algorithms of both types). Like NH, REMoDNaV is a *velocity-based* event classification algorithm. The algorithm comprises two major steps: preprocessing and event classification. The following sections detail individual analysis steps. For each step relevant algorithm parameters are given in parenthesis. Figure 1 provides an overview of the algorithm’s main components. Table 1 summarizes all parameters, and lists their default values. The computational definitions of the different eye movements (Hessels et al., 2018) are given within the event classification description. Note, however, that some of the computational definitions of eye movements can be adjusted to comply to alternative definitions by changing the algorithms’ parameters.

**Table 1:**
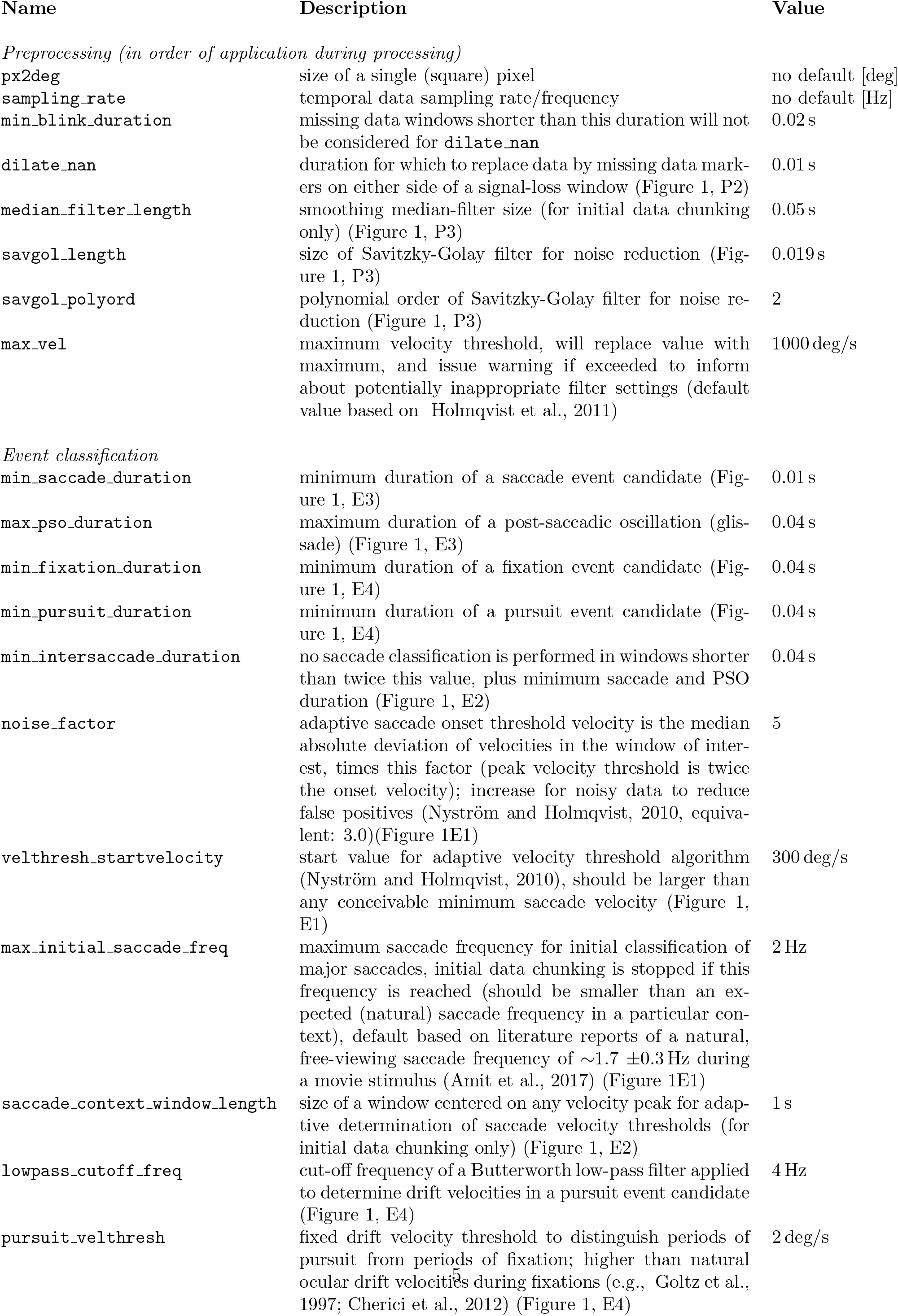
Exhaustive list of algorithm parameters, their default values, and units.

| Name | Description | Value |
| --- | --- | --- |
| *Preprocessing (in order of application during processing)* |  |  |
| px2deg | size of a single (square) pixel | no default [deg] |
| sampling_rate | temporal data sampling rate/frequency | no default [Hz] |
| min_blink_duration | missing data windows shorter than this duration will not be considered for `dilate_nan` | 0.02 s |
| dilate_nan | duration for which to replace data by missing data markers on either side of a signal-loss window (Figure 1, P2) | 0.01 s |
| median_filter_length | smoothing median-filter size (for initial data chunking only) (Figure 1, P3) | 0.05 s |
| savgol_length | size of Savitzky-Golay filter for noise reduction (Figure 1, P3) | 0.019 s |
| savgol_polyord | polynomial order of Savitzky-Golay filter for noise reduction (Figure 1, P3) | 2 |
| max_vel | maximum velocity threshold, will replace value with maximum, and issue warning if exceeded to inform about potentially inappropriate filter settings (default value based on Holmqvist et al., 2011) | 1000 deg/s |
| *Event classification* |  |  |
| min_saccade_duration | minimum duration of a saccade event candidate (Figure 1, E3) | 0.01 s |
| max_pso_duration | maximum duration of a post-saccadic oscillation (glissade) (Figure 1, E3) | 0.04 s |
| min_fixation_duration | minimum duration of a fixation event candidate (Figure 1, E4) | 0.04 s |
| min_pursuit_duration | minimum duration of a pursuit event candidate (Figure 1, E4) | 0.04 s |
| min_intersaccade_duration | no saccade classification is performed in windows shorter than twice this value, plus minimum saccade and PSO duration (Figure 1, E2) | 0.04 s |
| noise_factor | adaptive saccade onset threshold velocity is the median absolute deviation of velocities in the window of interest, times this factor (peak velocity threshold is twice the onset velocity); increase for noisy data to reduce false positives (Nyström and Holmqvist, 2010, equivalent: 3.0)(Figure 1E1) | 5 |
| velthresh_startvelocity | start value for adaptive velocity threshold algorithm (Nyström and Holmqvist, 2010), should be larger than any conceivable minimum saccade velocity (Figure 1, E1) | 300 deg/s |
| max_initial_saccade_freq | maximum saccade frequency for initial classification of major saccades, initial data chunking is stopped if this frequency is reached (should be smaller than an expected (natural) saccade frequency in a particular context), default based on literature reports of a natural, free-viewing saccade frequency of $\sim 1.7 \pm 0.3$ Hz during a movie stimulus (Amit et al., 2017) (Figure 1E1) | 2 Hz |
| saccade_context_window_length | size of a window centered on any velocity peak for adaptive determination of saccade velocity thresholds (for initial data chunking only) (Figure 1, E2) | 1 s |
| lowpass_cutoff_freq | cut-off frequency of a Butterworth low-pass filter applied to determine drift velocities in a pursuit event candidate (Figure 1, E4) | 4 Hz |
| pursuit_velthresh | fixed drift velocity threshold to distinguish periods of pursuit from periods of fixation; higher than natural ocular drift velocities during fixations (e.g., Goltz et al., 1997; Cherici et al., 2012) (Figure 1, E4) | 2 deg/s |

**Figure 1:**
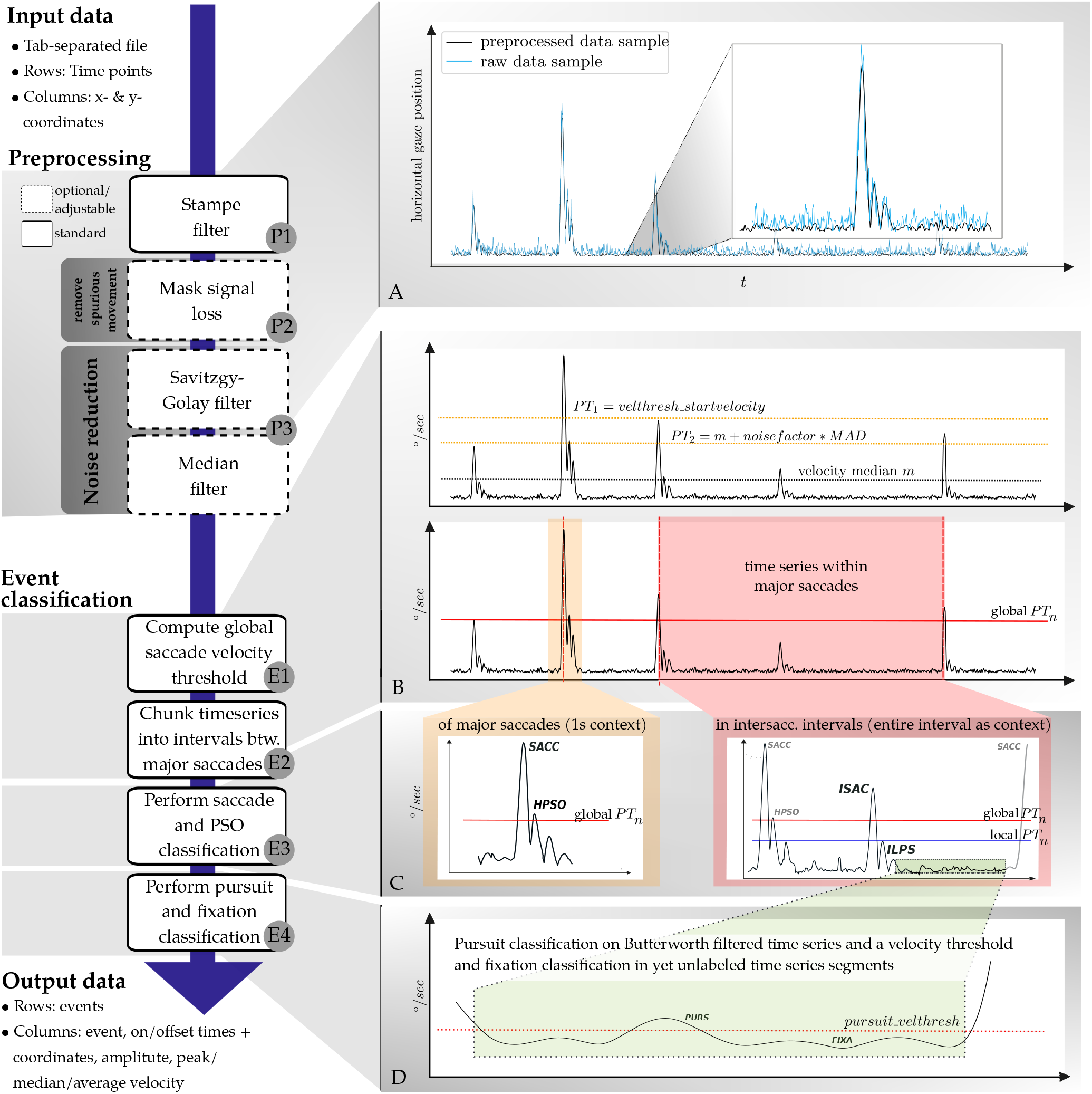
Schematic algorithm overview. (A) Preprocessing. The two plots show raw (blue) and processed (black) time series after preprocessing with the default parameter values (see Table 1 for details). (B) Adaptive saccade velocity computation and time series chunking. Starting from an initial velocity threshold (`velthresh_startvelocity`), a global velocity threshold is iteratively determined. The time series is chunked into intervals between the fastest saccades across the complete recording. (C) Saccade and PSO classification. Saccade on- and offsets, and PSO on- and offsets are classified based on adaptive velocity thresholds computed within the respective event contexts. The default context is either 1 s centered on the peak velocity for saccadic events used for time series chunking, or the entire time series chunk for intersaccadic intervals. PSOs are classified into low- or high-velocity PSOs depending on whether they exceed the saccade onset- or peak-velocity threshold. (D) Fixation and pursuit classification. Remaining unlabeled segments are filtered with a low-pass Butterworth filter. Samples exceeding a configurable pursuit velocity threshold (`pursuit_velthresh`) are classified as pursuits, and segments that do not qualify as pursuits are classified as fixations.

### Preprocessing

The goal of data preprocessing is to compute a time series of eye movement velocities on which the event classification algorithm can be executed, while jointly reducing non-eyemovement-related noise in the data as much as possible.

First, implausible spikes in the coordinate time series are removed with a heuristic spike filter (Stampe, 1993) (Figure 1, P1). This filter is standard in many eye tracking toolboxes and often used for preprocessing (e.g., Friedman et al., 2018). Data samples around signal loss (e.g., eye blinks) can be set to non-numeric values (NaN) in order to eliminate spurious movement signals without shortening the time series (`dilate_nan, min_blink_duration`; Figure 1, P2). This is motivated by the fact that blinks can produce artifacts in the eye-tracking signal when the eyelid closes and re-opens (Choe et al., 2016). Coordinate time series are temporally filtered in two different ways Figure 1, P3). A relatively large median filter (`median_filter_length`) is used to emphasize large amplitude saccades. This type of filtered data is later used for a coarse segmentation of a time series into shorter intervals between major saccades. Separately, data are also smoothed with a Savitzky-Golay filter (`savgol_{length,polyord}`). All event classification beyond the localization of major saccades for time series chunking is performed on this type of filtered data.

After spike-removal and temporal filtering, movement velocities are computed. To disregard biologically implausible measurements, a configurable maximum velocity (`max_vel`) is enforced—any samples exceeding this threshold are replaced by this set value.

### Event classification

#### Saccade velocity threshold

Except for a few modifications, REMoDNaV employs the adaptive saccade classification algorithm proposed by Nyström and Holmqvist (2010), where saccades are initially located by thresholding the velocity time series by a critical value. Starting from an initial velocity threshold (`velthresh_startvelocity`, termed *PT*_1_ in NH), the critical value is determined adaptively by computing the variance of sub-threshold velocities (*V*), and placing the new velocity threshold at:

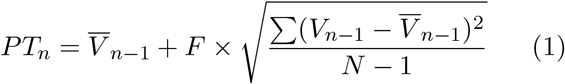

where *F* determines how many standard deviations above the average velocity the new threshold is located. This procedure is repeated until it stabilizes on a threshold velocity.

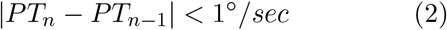

REMoDNaV alters this algorithm by using robust statistics that are more suitable for the non-normal distribution of velocities (Friedman et al., 2018), such that the new threshold is computed by:

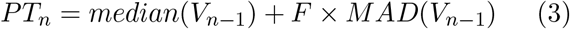

where *MAD* is the median absolute deviation, and *F* is a scalar parameter of the algorithm. This iterative process is illustrated in Figure 1, E1 (upper panel).

#### Time series chunking

As the algorithm aims to be applicable to prolonged recordings with potentially inhomogeneous noise levels, the time series needs to be split into shorter chunks to prevent the negative impact of sporadic noise flares on the aforementioned adaptive velocity thresholding procedure.

REMoDNaV implements this time-series chunking by determining a critical velocity on a median-filtered (`median_filter_length`) time series comprising the full duration of a recording (Figure 1, E2). Due to potentially elevated noise levels, the resulting threshold tends to overestimate an optimal threshold. Consequently, only periods of fastest eye movements will exceed this threshold. All such periods of consecutive above-threshold velocities are weighted by the sum of these velocities. Boundaries of time series chunks are determined by selecting such events sequentially (starting with the largest sums), until a maximum average frequency across the whole time series is reached (`max_initial_saccade_freq`). The resulting chunks represent data intervals between saccades of maximum magnitude in the respective data. Figure 1, E3 (right) exemplifies event classification within such an intersaccadic interval.

#### Classification of saccades and postsaccadic oscillations

Classification of these event types is identical to the NH algorithm, only the data context and metrics for determining the velocity thresholds differ. For saccades that also represent time series chunk boundaries (event label `SACC`), a context of 1 s (`saccade_context_window_length`) centered on the peak velocity is used by default, for any other saccade (event label `ISAC`) the entire time series chunk represents that context (Figure 1, E3).

Peak velocity threshold and on/offset velocity threshold are then determined by equation 3 with *F* set to 2 × `noise_factor` and `noise_factor`, respectively. Starting from a velocity peak, the immediately preceding and the following velocity minima that do not exceed the on/offset threshold are located and used as event boundaries. Qualifying events are rejected if they do not exceed a configurable minimum duration or violate the set saccade maximum proximity criterion (`min_saccade_duration, min_intersaccade_duration`).

As in NH, post-saccadic oscillations are events that immediately follow a saccade, where the velocity exceeds the saccade velocity threshold within a short time window (`max_pso_duration`). REMoD-NaV distinguishes low-velocity (event label LPSO for chunk boundary event, `ILPS` otherwise) and high-velocity oscillations (event label `HPSO` or `IHPS`), where the velocity exceeds the saccade onset or peak velocity threshold, respectively.

#### Pursuit and fixation classification

For all remaining, unlabeled time series segments that are longer than a minimum duration (`min_fixation_duration`), velocities are low-pass filtered (Butterworth, `lowpass_cutoff_freq`). Any segments exceeding a minimum velocity thresh-old (`pursuit_velthresh`) are classified as pursuit (event label `PURS`). Pursuit on/offset classification uses the same approach as that for saccades: search for local minima preceding and following the above threshold velocities. Any remaining segment that does not qualify as a pursuit event is classified as a fixation (event label `FIXA`) (Figure 1, E4).

#### Operation

REMoDNaV is free and open-source software, written in the Python language and released under the terms of the MIT license. In addition to the Python standard library it requires the Python packages NumPy (Oliphant, 2006), Matplotlib (Hunter, 2007), statsmodels (Seabold and Perktold, 2010), and SciPy (Jones et al., 2001-) as software dependencies. Furthermore, DataLad (Halchenko et al., 2013-), and Pandas (McKinney et al., 2010) have to be available to run the test battery. REMoDNaV itself, and all software dependencies are available on all major operating systems. There are no particular hardware requirements for running the software other than sufficient memory to load and process the data.

A typical program invocation looks like

~~~
remodnav <inputfile> <outputfile> \ <px2deg> <samplingrate>
~~~

where `<inputfile>` is the name of a tab-separated-value (TSV) text file with one gaze coordinate sample per line. An input file can have any number of columns, only the first two columns are read and interpreted as *X* and *Y* coordinates. Note that this constrains input data to a dense data representation, i.e. either data from eye trackers with fixed sampling frequency throughout the recording, or sparse data that has been transformed into a dense representation beforehand. The second argument `<outputfile>` is the file name of a BIDS-compliant (Gorgolewski et al., 2016) TSV text file that will contain a report on one classified eye movement event per line, with onset and offset time, onset and offset coordinates, amplitude, peak velocity, median velocity and average velocity. The remaining arguments are the only two mandatory parameters: the conversion factor from pixels to visual degrees, i.e., the visual angle of a single (square) pixel (`<px2deg>` in deg), and the temporal sampling rate (`<sampling_rate>` in Hz). Any other supported parameter can be added to the program invocation to override the default values.

A complete list of supported parameters (sorted by algorithm step) with their description and default value, are listed in Table 1. While the required user input is kept minimal, the number of configurable parameters is purposefully large to facilitate optimal parameterization for data with specific properties. Besides the list of classified events, a visualization of the classification results, together with a time course of horizontal and vertical gaze position, and velocities is provided for illustration and initial quality assessment of algorithm performance on each input data file.

### Validation analyses

The selection of datasets and analyses for validating algorithm performance was guided by three objectives: 1) compare to other existing solutions; 2) demonstrate plausible results on data from prolonged gaze coordinate recordings during viewing of dynamic, feature-rich stimuli; and 3) illustrate result robustness on lower-quality data. The following three sections each introduce a dataset and present the validation results for these objectives. All analysis presented here are performed using default parameters (Table 1), with no dataset-specific tuning other than the built-in adaptive behavior.

#### Algorithm comparison

Presently, Andersson et al. (2017) represents the most comprehensive comparative study on eye movement classification algorithms. Moreover, the dataset employed in that study was made publicly available. Consequently, evaluating REMoDNaV performance on these data and using their metrics offers a straightforward approach to relate this new development to alternative solutions.

The dataset provided by Andersson et al. (2017)^1^ consists of monocular eye gaze data produced from viewing stimuli from three distinct categories—images, moving dots and videos. The data release contains gaze coordinate time series (500 Hz sampling rate), and metadata on stimulus size and viewing distance. Importantly, each time point was manually classified by two expert human raters as one of six event categories: fixation, saccade, PSO, smooth pursuit, blink and undefined (a sample that did not fit any other category). A minor labeling mistake reported in Zemblys et al. (2018) was fixed prior to this validation analysis.

For each stimulus category, we computed the proportion of misclassifications per event type, comparing REMoDNaV to each of the human coders, and, as a baseline measure, the human coders against each other. A time point was counted as misclassified if the two compared classifications did not assign the same label. We limited this analysis to all time points that were labeled as fixation, saccade, PSO, or pursuit by any method, hence ignoring the rarely used NaN/blinks or “undefined” category. For a direct comparison with the results in Andersson et al. (2017), the analysis was repeated while also excluding samples labeled as pursuit. In the labeled data, there was no distinction made between high- and low-velocity PSOs, potentially because the literature following Nyström and Holmqvist (2010) did not adopt their differentiation of PSOs into velocity categories. All high- and low-velocity PSOs classified by REMoDNaV were therefore collapsed into a single PSO category. Table 2 shows the misclassification rates for all pair-wise comparisons, in all stimulus types. In comparison to the NH algorithm, after which the proposed work was modelled, REMoDNaV performed consistently better (32/93/70% average misclassification for NH, vs. 6.5/10.8/ 9.1% worst misclassification for REMoDNaV in categories images, dots, and videos). Compared to all ten algorithms evaluated in Andersson et al. (2017), REMoDNaV exhibits the lowest misclassification rates across all stimulus categories. When taking smooth pursuit events into account, the misclassification rate naturally increases, but remains comparably low. Importantly, it still exceeds the performance of all algorithms tested in Andersson et al. (2017) in the dots and video category, and performs among the best in the images category. Additionally, both with and without smooth pursuit, REMoDNaVs performance exceeds also that of a recent deep neural network trained specifically on video clips (Startsev et al., 2018, compare Table 7: 34% misclassification versus 31.5% for REMoDNaV).

**Table 2:** Proportion of samples in each stimulus category classified in disagreement between human coders (MN, RA) and the REMoDNaV algorithm (AL). The MC (misclassification) column lists proportions considering all four event categories (fixation, saccade, PSO, pursuit), while the w/oP (without pursuit) column excludes pursuit events for a direct comparison with Andersson et al. (2017, Tables 8-10). The remaining columns show the percentage of labels assigned to incongruent time points by each rater (deviation of their sum from 100% is due to rounding).

| **Images** |  |  |  |  |  |  |  |
| --- | --- | --- | --- | --- | --- | --- | --- |
| Comp | MC | w/oP | Coder | Fix | Sac | PSO | SP |
| MN-RA | 6.1 | 3.0 | MN | 70 | 9 | 21 | 0 |
| — | — | — | RA | 13 | 15 | 20 | 53 |
| MN-AL | 23.1 | 6.5 | MN | 86 | 2 | 11 | 2 |
| — | — | — | AL | 5 | 13 | 6 | 75 |
| RA-AL | 22.8 | 6.4 | RA | 77 | 3 | 11 | 9 |
| — | — | — | AL | 13 | 13 | 6 | 68 |
| **Dots** |  |  |  |  |  |  |  |
| Comp | MC | w/oP | Coder | Fix | Sac | PSO | SP |
| MN-RA | 10.7 | 4.2 | MN | 11 | 10 | 9 | 71 |
| — | — | — | RA | 64 | 7 | 6 | 23 |
| MN-AL | 18.6 | 8.2 | MN | 9 | 5 | 8 | 78 |
| — | — | — | AL | 77 | 6 | 2 | 15 |
| RA-AL | 22.8 | 10.8 | RA | 28 | 4 | 6 | 61 |
| — | — | — | AL | 59 | 7 | 2 | 31 |
| **Videos** |  |  |  |  |  |  |  |
| Comp | MC | w/oP | Coder | Fix | Sac | PSO | SP |
| MN-RA | 18.5 | 4.0 | MN | 75 | 3 | 8 | 15 |
| — | — | — | RA | 16 | 4 | 3 | 77 |
| MN-AL | 31.5 | 7.9 | MN | 57 | 1 | 6 | 36 |
| — | — | — | AL | 36 | 5 | 3 | 55 |
| RA-AL | 28.5 | 9.1 | RA | 38 | 3 | 5 | 55 |
| — | — | — | AL | 53 | 6 | 5 | 35 |

Figure 2 shows confusion patterns for a comparison of algorithm classifications with human labeling and displays the similarity between classification decisions with Jaccard indices (JI; Jaccard, 1901). The JI is bound in range [0, 1] with higher values indicating higher similarity. A value of 0.93 in the upper left cell of the very first matrix in Figure 2 for example indicates that 93% of frames that are labeled as a fixation by human coders RA and MN are the same. This index allows to quantify the similarity of classifications independent of values in other cells. While REMoDNaV does not achieve a labeling similarity that reaches the human inter-rater agreement, it still performs well. In particular, the relative magnitude of agreement with each individual human coder for fixations, saccades, and PSOs, resembles the agreement between the human coders. Classification of smooth pursuits is consistent with human labels for the categories moving dots, and videos. However, there is a substantial confusion of fixation and pursuit for the static images. In a real-world application of REMoDNaV, pursuit classification could be disabled (by setting a high pursuit velocity threshold) for data from static images, if the occurrence of pursuit events can be ruled out a priori. For this evaluation, however, no such intervention was made.

**Figure 2:**
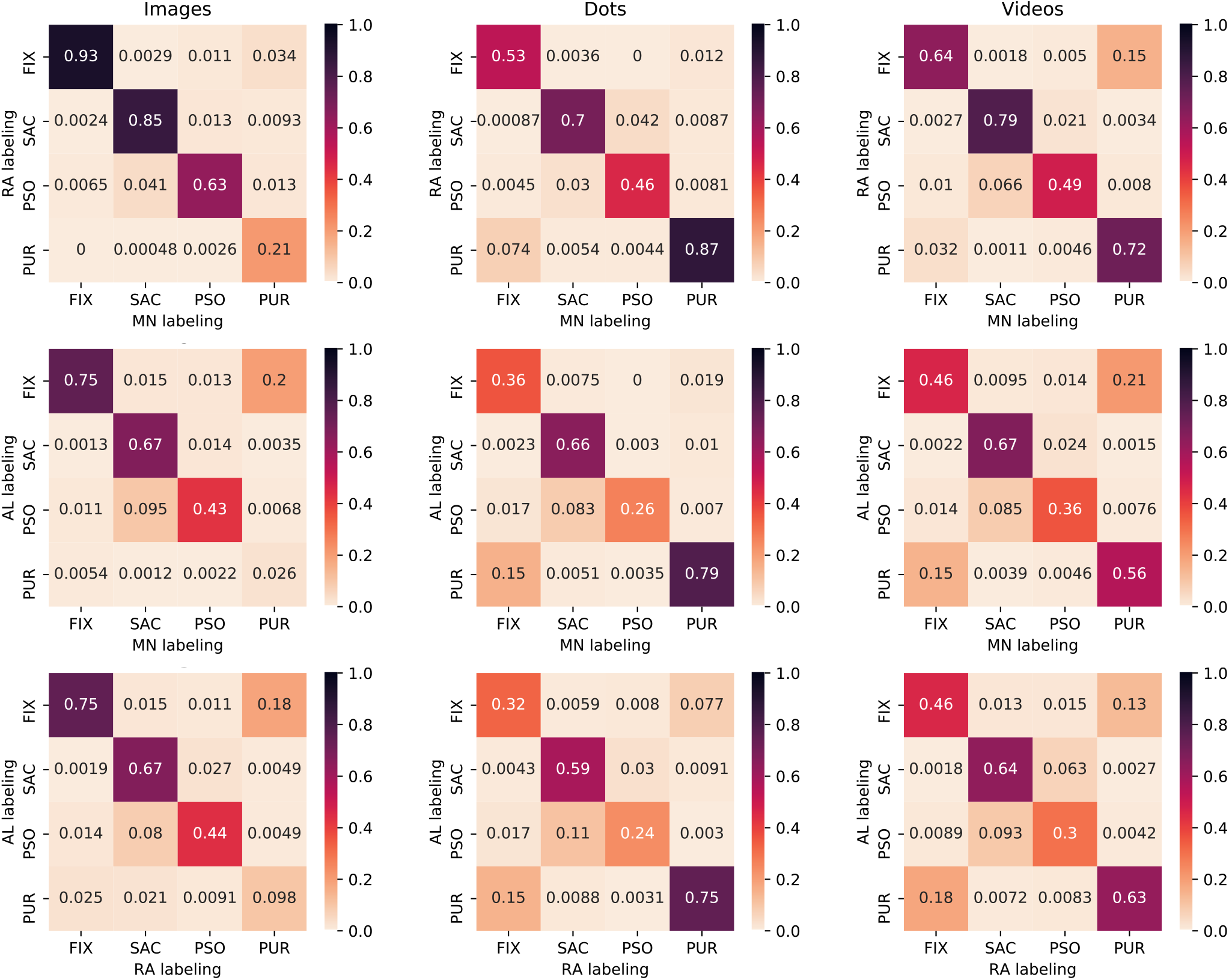
Confusion patterns for pairwise eye movement classification comparison of both human raters (MN and RA; Andersson et al., 2017) and the REMoDNaV algorithm (AL) for gaze recordings from stimulation with static images (left column), moving dots (middle column), and video clips (right column). All matrices present gaze sample based Jaccard indices (JI; Jaccard, 1901). Consequently, the diagonals depict the fraction of time points labeled congruently by both raters in relation to the number of timepoints assigned to a particular event category by any rater.

In addition to the confusion analysis and again following Andersson et al. (2017), we computed Cohen’s Kappa (Cohen, 1960) for an additional measure of similarity between human and algorithm performance. It quantifies the sample-by-sample agreement between two ratings following equation 4:

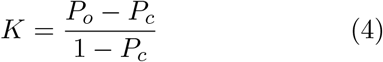

where *P*_*o*_ is the observed proportion of agreement between the ratings, and *P*_*c*_ is the proportion of chance agreement. A value of *K* = 1 indicates perfect agreement, and *K* = 0 indicates chance level agreement. Table 3 displays the resulting values between the two human experts, and REMoDNaV with each of the experts, for each stimulus category and the three event types used in Andersson et al. (2017), namely fixations, saccades, and PSOs (compare to Andersson et al. (2017), table 7). For all event types and stimulus categories, REMoDNaV performs on par or better than the original NH algorithm, and in many cases on par or better than the best of all algorithms evaluated in Andersson et al. (2017) within an event or stimulus type.

**Table 3:** Cohen’s Kappa reliability between human coders (MN, RA), and REMoDNaV (AL) with each of the human coders.

| **Fixations** |  |  |  |
| --- | --- | --- | --- |
| Comparison | Images | Dots | Videos |
| MN versus RA | 0.84 | 0.65 | 0.65 |
| AL versus RA | 0.55 | 0.37 | 0.44 |
| AL versus MN | 0.52 | 0.45 | 0.39 |
| **Saccades** |  |  |  |
| Comparison | Images | Dots | Videos |
| MN versus RA | 0.91 | 0.81 | 0.87 |
| AL versus RA | 0.78 | 0.72 | 0.76 |
| AL versus MN | 0.78 | 0.78 | 0.79 |
| **PSOs** |  |  |  |
| Comparison | Images | Dots | Videos |
| MN versus RA | 0.76 | 0.62 | 0.65 |
| AL versus RA | 0.59 | 0.38 | 0.45 |
| AL versus MN | 0.58 | 0.41 | 0.51 |

In order to further rank the performance of the proposed algorithm with respect to the ten algorithms studied in Andersson et al. (2017), we followed their approach to compute root mean square deviations (RMSD) from human labels for event duration distribution characteristics (mean and standard deviation of durations, plus number of events) for each stimulus category (images, dots, videos) and event type (fixations, saccades, PSOs, pursuits). This measure represents a scalar distribution dissimilarity score that can be used as an additional comparison metric of algorithm performance that focuses on overall number and durations of classified events, instead of sample-by-sample misclassification. The RMSD measure has a lower bound of 0.0 (identical to the average of both human raters), with higher values indicating larger differences (for detail information on the calculation of this metric see Andersson et al., 2017).

Table 4 is modelled after Andersson et al. (2017, Tables 3-6), appended with REMoDNaV, showing RMSD based on the scores of human raters given in the original tables. As acknowledged by the authors, the absolute value of the RMSD scores is not informative due to scaling with respect to the respective maximum value of each characteristic. Therefore, we converted RMSDs for each algorithm and event type into zero-based ranks (lower is more human-like).

**Table 4:**
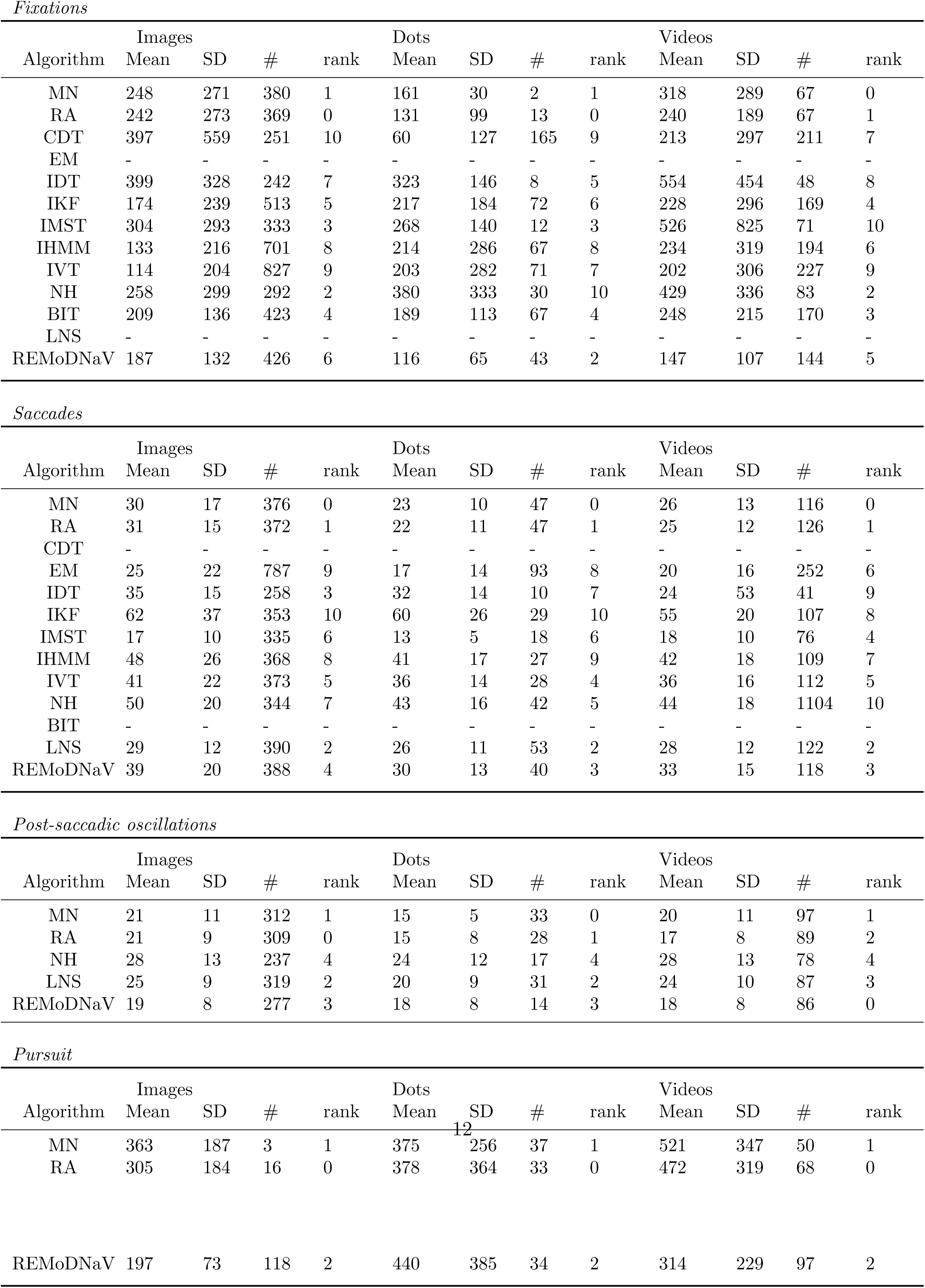
Comparison of event duration statistics (mean, standard deviation, and number of events) for image, dot, and video stimuli. This table is modeled after Andersson et al. (2017, Tables 3-6), and root-mean-square-deviations (RMSD) from human raters are shown for fixations, saccades, PSOs, and pursuit as zero-based ranks (rank zero is closest to the average of the two human raters). Summary statistics for all algorithms used in Andersson et al. (2017) were taken from their publicly available GitHub repository (github.com/richardandersson/EyeMovementDetectorEvaluation). Cohens Kappa was computed for the complete set of algorithms in Andersson et al. (2017) and REMoDNaV.

The LNS algorithm (Larsson et al., 2013) was found to have the most human-like performance for saccade and PSO classification in Andersson et al. (2017). REMoDNaV performs comparable to LNS for both event types (saccades: 2.0 vs. 3.3; PSOs: 2.3 vs. 2.0, mean rank across stimulus categories for LNS and REMoDNaV, respectively).

Depending on the stimulus type, different algorithms performed best for fixation classification. NH performed best for images and videos, but worst for moving dots. REMoDNaV outperforms all other algorithms in the dots category, and achieves rank 5 and 6 (middle range) for videos and images, respectively. Across all stimulus and event categories, REMoDNaV achieves a mean ranking of 2.9, and a mean ranking of 3.2 when not taking smooth pursuit into account.

Taken together, REMoDNaV yields classification results that are, on average, more human-like than any other algorithm tested on the dataset and metrics put forth by Andersson et al. (2017). In particular, its performance largely equals or exceeds that of the original NH algorithm. NH outperforms it only for fixation classification in the image and video category, but in these categories REMoDNaV also classifies comparatively well. These results are an indication that the changes to the NH algorithm proposed here to improve upon its robustness are not detrimental to its performance on data from conventional paradigms and stimuli.

### Prolonged viewing of dynamic stimuli

Given that REMoDNaV yielded plausible results for the “video” stimulus category data in the Andersson et al. (2017) dataset (Figure 2, and Table 4, right columns), we determined whether it is capable of analyzing data from dynamic stimulation in prolonged (15 min) recordings.

As a test dataset we used publicly available eye tracking data from the *studyforrest.org* project, where 15 participants were recorded watching a feature-length (≈2 h) movie in a laboratory setting (Hanke et al., 2016). Eye movements were measured by an Eyelink 1000 with a standard desktop mount (software version 4.51; SR Research Ltd., Mississauga, Ontario, Canada) and a sampling rate of 1000 Hz. The movie stimulus was presented on a 522 × 294 mm LCD monitor at a resolution of 1920×1280 px and a viewing distance of 85 cm. Participants watched the movie in eight approximately 15 min long segments, with recalibration of the eye tracker before every segment.

As no manual eye movement event labeling exists for these data, algorithm evaluation was limited to a comparison of marginal distributions and well-known properties, such as the log-log-linear relationship of saccade amplitude and saccade peak velocity (Bahill et al., 1975). Figure 3 (top row) depicts this main sequence relationship. Additionally, Figure 4 (top row) shows duration histograms for all four event types across all participants. Shapes and locations of these distributions match previous reports in the literature, such as a strong bias towards short (less than 500 ms) fixations for dynamic stimuli (Dorr et al., 2010, Fig. 3), peak number of PSOs with durations between 10-20 ms (Nyström and Holmqvist, 2010, Fig. 11), and a non-Gaussian saccade duration distribution located below 100 ms (Nyström and Holmqvist, 2010, Fig. 8, albeit for static scene perception). Overall, the presented summary statistics suggest that REMoDNaV is capable of classifying eye movements with plausible characteristics, in prolonged gaze recordings. A visualization of such a classification result is depicted in Figure 5 (top row).

**Figure 3:**
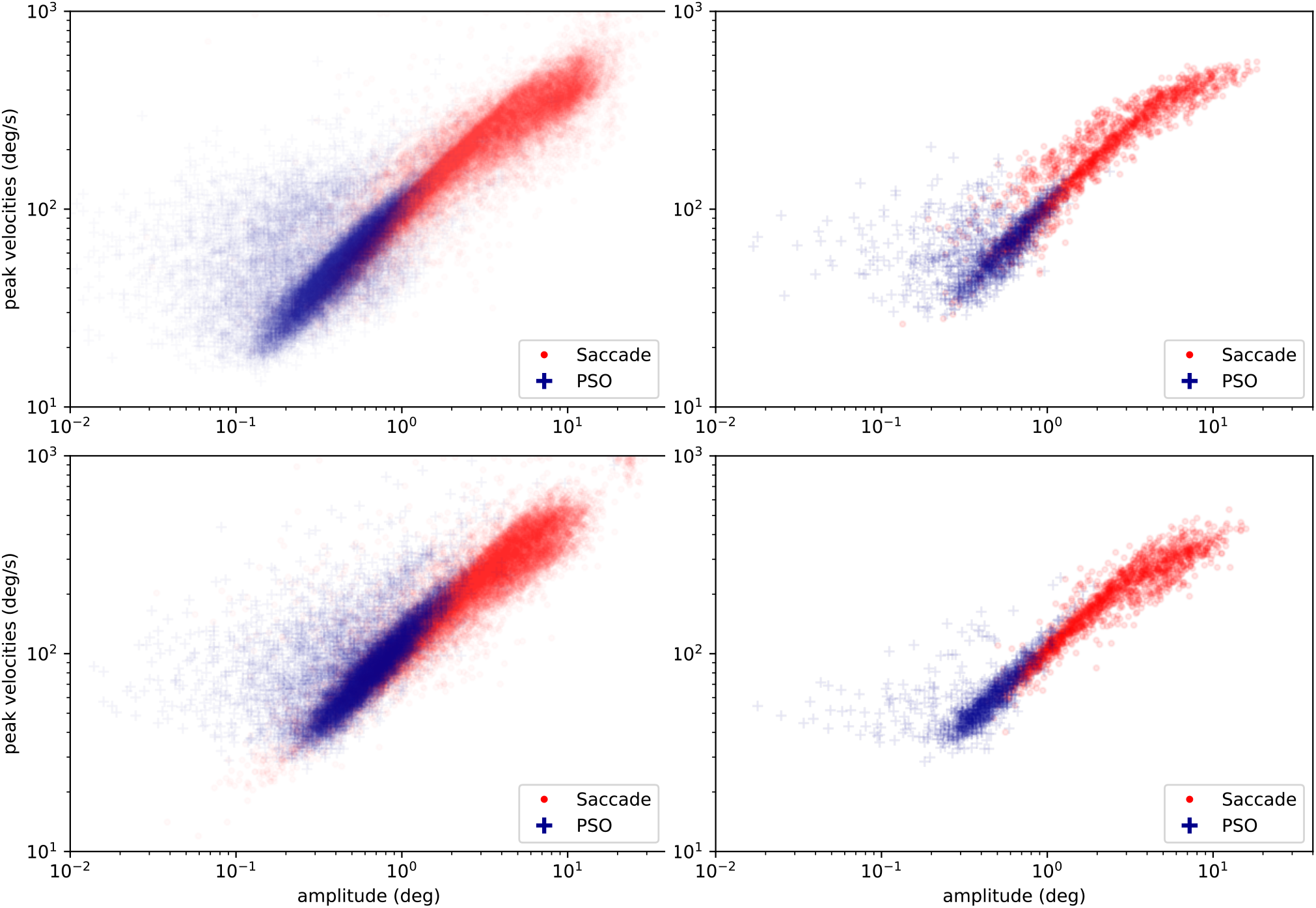
Main sequence of eye movement events during one 15 minute sequence of the movie (segment 2) for lab (top), and MRI participants (bottom). Data across all participants per dataset is shown on the left, and data for a single exemplary participant on the right.

**Figure 4:**
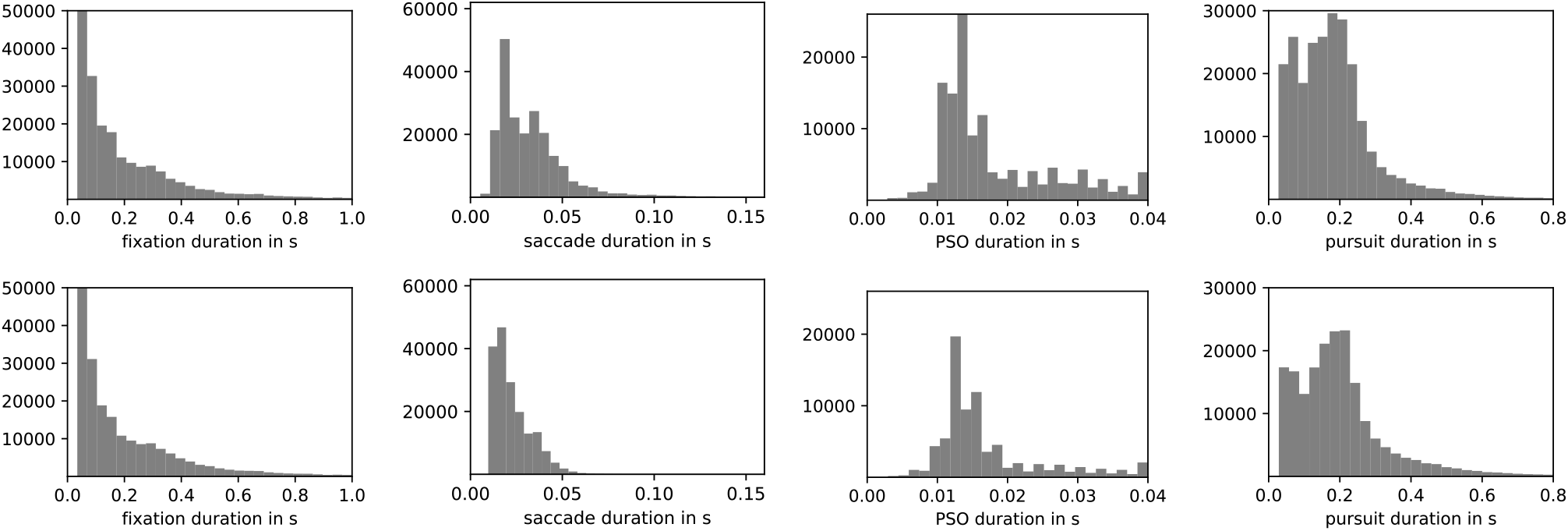
Comparison of eye movement event duration distributions for the high-quality lab sample (top row), and the lower quality MRI sample (bottom row) across all participants (each *N* = 15), and the entire duration of the same feature-length movie stimulus. All histograms depict absolute number of events. Visible differences are limited to an overall lower number of events, and fewer long saccades for the MRI sample. These are attributable to a higher noise level and more signal loss (compare Hanke et al., 2016, Fig. 4b) in the MRI sample, and to stimulus size differences (23.75° MRI vs. 34° lab).

**Figure 5:**
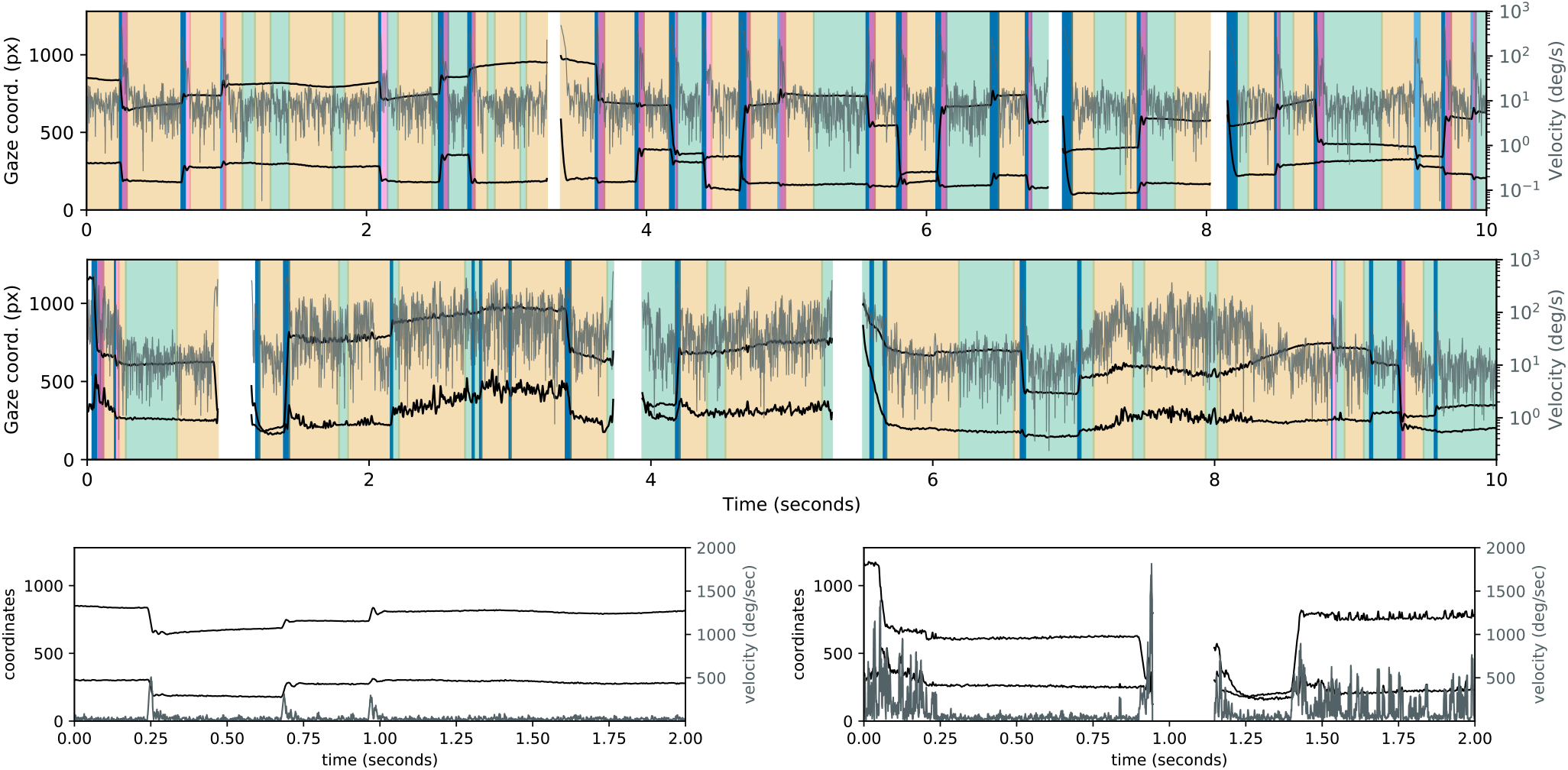
Exemplary eye movement classification results for the same 10 s excerpt of a movie stimulus for a single participant in the high quality lab sample (top), and in the lower quality MRI sample (middle). The plots show filtered gaze coordinates (black), computed velocity time series (gray) overlayed on the eye movement event segmentation with periods of fixation (green), pursuit (beige), saccades (blue), and high/low-velocity post-saccadic oscillations (dark/light purple). The bottom panel shows the first 2 s of unfiltered gaze coordinates (black) and unfiltered velocity time series (gray) for lab (left) and mri (right) sample in greater detail. The variable noise level, and prolonged signal loss (white in top panel) visible in the MRI sample represent a challenge for algorithms. REMoDNaV uses an adaptive approach that determines major saccade events first, and subsequently tunes the velocity threshold to short time windows between these events. Figures like this accompany the program output to facilitate quality control and discovery of inappropriate preprocessing and classification parameterization.

### Lower-quality data

An explicit goal for REMoDNaV development was robust performance on lower-quality data. While lack of quality cannot be arbitrarily compensated and can inevitably lead to misses in eye movement classification if too high, it is beneficial for any further analysis if operation on noisy data does not introduce unexpected event property biases.

In order to investigate noise-robustness we ran REMoDNaV on another publicly available dataset from the *studyforrest.org* project, where 15 different participants watched the exact same movie stimulus, but this time while lying on their back in the bore of an MRI scanner (Hanke et al., 2016). These data were recorded with a different Eyelink 1000 (software version 4.594) equipped with an MR-compatible telephoto lens and illumination kit (SR Research Ltd., Mississauga, Ontario, Canada) at 1000 Hz during simultaneous fMRI acquisition. The movie was presented at a viewing distance of 63 cm on a 26 cm (1280 × 1024 px) LCD screen in 720p resolution at full width, yielding a substantially smaller stimulus size, compared to the previous stimulation setup. The eye tracking camera was mounted outside the scanner bore and recorded the participants’ left eye at a distance of about 100 cm. Compared to the lab-setup, physical limitations of the scanner environment, and sub-optimal infrared illumination led to substantially noisier data, as evident from a larger spatial uncertainty (Hanke et al., 2016, Technical Validation), a generally higher amount of data loss, and more events with a velocity above 800 deg/s. Following common data quality criteria used to warrant exclusion by Holmqvist et al. (2012), a higher amount of zero values, a greater number of events with a velocity above 800 deg/s, and lower spatial accuracy can be indicative of lower quality data. The average amount of data loss in the MRI sample was three times higher than in the laboratory setting (15.1% versus 4.1% in the lab), with six out of 15 subjects having one or more movie segments with data loss greater than 30%. In the laboratory setting, in comparison, zero out of 15 subjects had one or more movie segments with data loss greater than 30% (Hanke et al., 2016, Table 1). Figure 6 highlights the higher amount of extreme velocities in the MRI sample, even though the stimulus size was smaller than in the laboratory setting. Finally, the average spatial accuracy at the start of a recording, assessed with a 13-point calibration procedure, was 0.58 degrees of visual angle for the MRI sample and 0.45 degrees for the lab sample (Hanke et al., 2016, Technical Validation). An example of the amplified and variable noise pattern is shown in Figure 5 (bottom row, gray lines). Except for the differences in stimulation setup, all other aspects of data acquisition, eye tracker calibration, and data processing were identical to the previous dataset.

**Figure 6:**
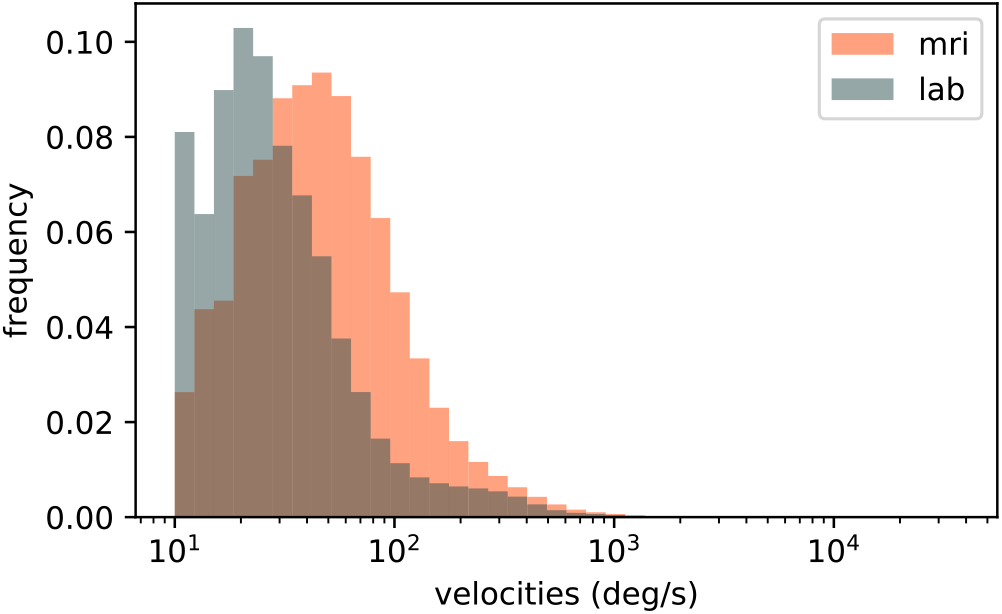
Comparison of sample velocity distributions for MRI and laboratory setting across all measurements and participants (excluding samples during periods of signal-loss). The MRI sample exhibits a larger fraction of higher velocities, despite a 30% smaller stimulus size.

We performed the identical analysis as before, in order to compare performance between a high and lower-quality data acquisition. This approach differs from the common approach of adding increasing levels of artificial noise to data (as done for example in Hessels et al. (2017)), but bears the important advantage of incorporating real lower-quality data characteristics instead of potentially inappropriate or unnatural noise. Figures 3-5 depict the results for the lab-quality dataset, and the MRI-scanner dataset in the top and bottom rows, respectively.

Overall, the classification results exhibit strong similarity, despite the potential behavioral impact of watching a movie while lying on their back and looking upwards on the participants, or the well known effect of increasing fatigue (Tagliazucchi and Laufs, 2014) during a two-hour session in an MRI-scanner. In particular, saccade amplitude and peak velocity exhibit a clear main-sequence relationship that resembles that found for the lab acquisition (Figure 3). Duration distributions for fixations, PSOs, and pursuits are strikingly similar between the two datasets (Figure 4), except for a generally lower number of classified events for the MRI experiment, which could be explained by the higher noise level and fraction of signal loss. There is a notable difference regarding the saccade duration distributions, with a bias towards shorter saccades in the MRI dataset. This effect may be attributable to the differences in stimulus size (30% smaller in the MRI environment).

## Conclusion

Based on the adaptive, velocity-based algorithm for fixation, saccade, and PSO classification by Nyström and Holmqvist (2010), we have developed an improved algorithm that performs robustly on prolonged or short recordings with dynamic stimulation, with potentially variable noise levels, and also supports the classification of smooth pursuit events. Through a series of validation analyses we have shown that its performance is comparable to or better than ten other contemporary algorithms, and that plausible classification results are achieved on high and lower quality data. These aspects of algorithm capabilities and performance suggest that REMoDNaV is a state-of-the-art tool for eye movement classification with particular relevance for emerging complex data collections paradigms with dynamic stimulation, such as the combination of eye tracking and functional MRI in simultaneous measurements.

The proposed algorithm is rule-based, hence can be applied to data without prior training, apart from the adaptive estimation of velocity thresholds. This aspect distinguishes it from other recent developments based on deep neural networks (Startsev et al., 2018), and machine-learning in general (Zemblys et al., 2018). Some statistical learning algorithms require (labeled) training data, which can be a limitation in the context of a research study. However, in its present form REMoDNaV cannot be used for real-time data analysis, as its approach for time series chunking is based on an initial sorting of major saccade events across the entire time series. The proposed algorithm presently does not support the classification of eye blinks as a category distinct from periods of general signal loss. While such a feature could potentially be added, the current default preprocessing aims at removing blink-related signal. The algorithm maintains a distinction between high- and low-velocity PSOs first introduced by Nyström and Holmqvist (2010), although, to our knowledge, the present literature does not make use of such a distinction. Algorithm users are encouraged to decide on a case-by-case basis whether to lump these event categories together into a general PSO category, as done in our own validation analyses. As a general remark it is also noteworthy that eye tracking systems using pupil corneal reflection (pupil-CR) eye tracking may bias data towards premature PSO onset times and inflated PSO peak velocities (see Hooge et al. (2016)). In deciding whether and how to interpret PSO events, it needs to be considered whether the eye tracking device may have introduced biases in the data. Lastly, the evaluation results presented here are based on data with a relatively high temporal resolution (0.5 and 1 kHz). While the algorithm does not impose any hard constraints on data acquisition parameters, its performance on data from low-end, consumer grade hardware (e.g., 50 Hz sampling rate) has not been tested.

Just as Andersson et al. (2017), we considered human raters as a gold standard reference for event classification when evaluating algorithms. The implications of the results presented herein are hence only valid if this assumption is warranted. Some authors voice concerns (e.g., Komogortsev et al., 2010), regarding potential biases that may limit generalizability. Nevertheless, human-made event labels are a critical component of algorithm validation, as pointed out by Hooge et al. (2018).

The validation analyses presented here are based on three different datasets: a manually annoted dataset (Andersson et al., 2017), and two datasets with prolonged recordings using movie stimuli (Hanke et al., 2016). Beyond our own validation, a recent evaluation of nine different smooth pursuit algorithms by Startsev, Agtzidis and Dorr as part of their recent paper (Startsev et al., 2018) also provides metrics for REMoDNaV. In their analysis, algorithm performance was evaluated against a partially hand-labelled eye movement annotation of the Hollywood2 dataset (Mathe and Sminchisescu, 2012). We refrain from restating their methodology or interpreting their results here, but encourage readers to consult this independent report^2^.

REMoDNaV aims to be a readily usable tool, available as cross platform compatible, free and open source software, with a simple command line interface and carefully chosen default settings. However, as evident from numerous algorithm evaluations (e.g., Andersson et al., 2017; Larsson et al., 2013; Zemblys et al., 2018; Komogortsev et al., 2010), different underlying stimulation, and data characteristics can make certain algorithms or parameterizations more suitable than others for particular applications. The provided implementation of the REMoDNaV algorithm (Hanke et al., 2019) acknowledges this fact by exposing a range of parameters through its user interface that can be altered in order to tune the classification for a particular use case.

The latest version of REMoDNaV can be installed from PyPi^3^ via pip install remodnav. The source code of the software can be found on Github^4^. All reports on defects and enhancement can be submitted there. The analysis code underlying all results and figures presented in this paper, as well as the LATEX sources, are located in another Github repository^5^. All required input data, from Andersson et al. (2017) and the *studyforrest.org* project, are referenced in this repository at precise versions as DataLad^6^ subdatasets, and can be obtained on demand. The repository constitutes an automatically reproducible research object, and readers interested in verifying the results and claims of our paper can recompute and plot all results with a single command after cloning the repository.

## Author contributions

AD, MH conceived and implemented the algorithm. AD, AW, MH validated algorithm performance. AD, AW, MH wrote the manuscript.

## Footnotes

1 github.com/richardandersson/EyeMovementDetectorEvaluation

2 https://www.michaeldorr.de/smoothpursuit/

3 https://pypi.org/project/remodnav

4 https://github.com/psychoinformatics-de/remodnav

5 https://github.com/psychoinformatics-de/paper-remodnav/

6 http://datalad.org

